# Mitochondrial ubiquinone-mediated longevity is marked by reduced cytoplasmic protein translation

**DOI:** 10.1101/308858

**Authors:** Marte Molenaars, Georges E. Janssens, Toon Santermans, Marco Lezzerini, Rob Jelier, Alyson W. MacInnes, Riekelt H. Houtkooper

## Abstract

Mutations in the *clk-1* gene impair mitochondrial ubiquinone biosynthesis and extend the lifespan of *C. elegans.* We demonstrate here that this life extension is linked to the repression of cytoplasmic protein translation. *Clk-1* mutations inhibit polyribosome formation similarly to *daf-2* mutations that dampen insulin signaling. Comparisons of total versus polysomal RNAs in *clk-1* mutants reveal a reduction in the translational efficiencies of mRNAs coding for elements of the translation machinery and an increase in those coding for the oxidative phosphorylation and autophagy pathways. Knocking down the transcription initiation factor TAF-4, a protein that becomes sequestered in the cytoplasm during early embryogenesis to induce transcriptional silencing, ameliorates the *clk-1* inhibition of polyribosome formation. These results underscore a prominent role for the repression of cytoplasmic protein translation in eukaryotic lifespan extension, and suggest that mutations impairing mitochondrial function are able to exploit this repression similarly to reductions of insulin signaling. Moreover, this report reveals an unexpected role for TAF-4 as a repressor of polyribosome formation when ubiquinone biosynthesis is compromised.

## Introduction

Slowing down mitochondrial metabolism is a well-known means of increasing the lifespan of multiple species (Breitenbach et al., 2014, Copeland et al., 2009, Dillin et al., 2002, Houtkooper et al., 2013). In *C. elegans*, one such model is the *clk-1(mq30)* mutant that harbors a deletion in the gene encoding a ubiquinone (UQ) biosynthesis enzyme (Felkai et al., 1999). Ubiquinone is a redox-active lipid that plays a central role in the electron transport chain and mitochondrial oxidative phosphorylation (OXPHOS) by carrying electrons from complexes I and II to complex III (Stefely & Pagliarini, 2017). In addition to lifespan extension and decreased respiration, *clk-1* mutants are developmentally delayed and have reduced progeny (Felkai et al., 1999). Mammalian *Clk1*, known as Mclk-1 or Coq7, is also involved in lifespan (Liu et al., 2005, Stepanyan et al., 2006, Takahashi et al., 2014). Transgenic mice, rescued from embryonic lethality via the transgenic expression of mouse clk-1, with reduced *Clk1* expression live longer and have smaller bodies than wild-type mice, demonstrating a conserved role for *clk-1* in longevity across species (Takahashi et al., 2014). However it is not fully understood how the reduction of ubiquinone biosynthesis extends the lifespan of these animals.

Until recently, CLK-1 protein was thought to reside exclusively in the mitochondria. However, a distinct nuclear form of CLK-1 has been reported to independently regulate lifespan by mitochondrial-nuclear retrograde signaling (Monaghan et al., 2015). Adding back a copy of the *clk-1* gene that lacks the mitochondrial targeting signal (MTS) to the *clk-1* deletion mutant led to a subtle but significant reduction in the lifespan extension phenotype observed in the full *clk-1* deletion mutant (Monaghan et al., 2015). However, the existence and function of a nuclear form of CLK-1 remains controversial. Other studies fail to detect any nuclear localization of the MCLK1/CLK-1 proteins or any biological activity of a *C. elegans* CLK-1 protein devoid of an MTS (Liu et al., 2017).

The importance of communication between stressed mitochondria and the rest of the cell is becoming increasingly appreciated (Quiros et al., 2016). One such reaction to stress is carried out by the mitochondrial unfolded protein response (UPR^mt^) (Baker et al., 2012, Durieux et al., 2011, Houtkooper et al., 2013). GCN-2, an eIF2α kinase that modulates cytosolic protein synthesis, is phosphorylated in response to the activated UPR^mt^ in *clk-1* mutants in a manner required for lifespan extension (Baker et al., 2012). Another non-mitochondrial factor, TATA-binding protein associated factor (TAF-4), is a transcription factor that has been implicated in the extension of *clk-1* mutant lifespan (Khan et al., 2013). TAF-4 was identified in an RNA interference (RNAi) screen for transcription factors required for the lifespan extension of *clk-1* mutants (Khan et al., 2013). Beyond *clk-1*, TAF-4 was also required for the lifespan extension phenotype of two mutants of the mitochondrial electron transport chain (*isp-1* and tpk-1) (Khan et al., 2013). TAF-4 is a component of the TFIID mRNA transcription complex and is best known for its role in transcriptional silencing in early embryogenesis by becoming sequestered in the cytoplasm due to phosphorylation by OMA-1 (Walker et al., 2001). The loss of TAF-4 in *C. elegans* suppresses the lifespan extension phenotype induced by *clk-1* mutations, however the mechanism through which this happens is not known.

Reducing protein translation is another well-known means by which eukaryotes extend lifespan (Hansen et al., 2007, Pan et al., 2007). Caloric restriction, amino acid reduction, and knocking down myriad factors involved in protein translation (such as ribosomal proteins, elongation/initiation factors, or tRNA synthetases) all increase lifespan (Hansen et al., 2007, Masoro, 2000, Min & Tatar, 2006, Pan et al., 2007). The genetic or pharmacological inhibition of the mTOR/nutrient sensing pathway or insulin receptor signaling pathway is also marked by reduced protein translation rates and a suppression of polyribosome formation (Genolet et al., 2008, Stout et al., 2013). Proteomic analysis of the *daf-2* mutants previously revealed a substantial reduction of ribosomal proteins, translation factors, and protein metabolism components coupled to a repression of polyribosome formation (Stout et al., 2013). Moreover, our previous work demonstrated that similarly to *daf-2* mutants, mutations in *clk-1* worms also result in a strong repression of polyribosome formation (Essers et al., 2015). This led us to hypothesize the existence of unexplored regulatory links between dysfunctional mitochondria and cytoplasmic protein translation that are contributing to eukaryotic lifespan extension. In particular, we investigated which subsets of mRNAs are preferentially translated in the longer-lived *C. elegans* with impaired ubiquinone biosynthesis.

## Results

### RNAseq of *clk-1* strains shows major changes in transcription

To uncover how the transcriptome of *C. elegans* changes on deletion of *clk-1* and the influence of the *clk-1* mitochondrial localization signal, we performed next-generation sequencing of total RNAs (depleted of rRNA) isolated from worms at the L4 stage that precedes young adulthood. Total RNA pools depleted of rRNA isolated from *clk-1(qm30)* mutants were compared with (1) *clk-1(qm30)^+WT^* mutants rescued with the wildtype *clk-1* and (2) *clk-1(qm30)^+nuc^* mutants with predominantly the nuclear form of *clk-1* (Fig 1A) that was described in (Monaghan et al., 2015). The *clk-1(qm30)^+WT^* mutants have wild-type phenotypes and are rescued with fully functioning *clk-1* and, while the *clk-1(qm30)^+nuc^* mutants have *clk-1* only residing in the nucleus and not in the mitochondria (Monaghan et al., 2015). The principal component analysis (PCA) plot and correlation matrix show the expected clustering of the biological triplicates of each strain (Fig 1B-C). Major changes were observed in the RNA pool of the *clk-1(mq30)^+WT^* mutants compared with either *clk-1(mq30)* or *clk-1(mq30)^+nuc^*, with 9626 and 8685 genes being differentially expressed, respectively. Far fewer differentially expressed genes were measured when comparing *clk-1(mq30)* to *clk-1(mq30)^+nuc^*, with only 247 genes upregulated and 571 genes downregulated. The subtle changes in RNA expression are consistent with the observation that introducing the nuclear form of *clk-1* in *clk-1(mq30)* results in only small phenotypic changes compared to the *clk-1(mq30)* mutants (Liu et al., 2017, Monaghan et al., 2015).

**Figure 1.**
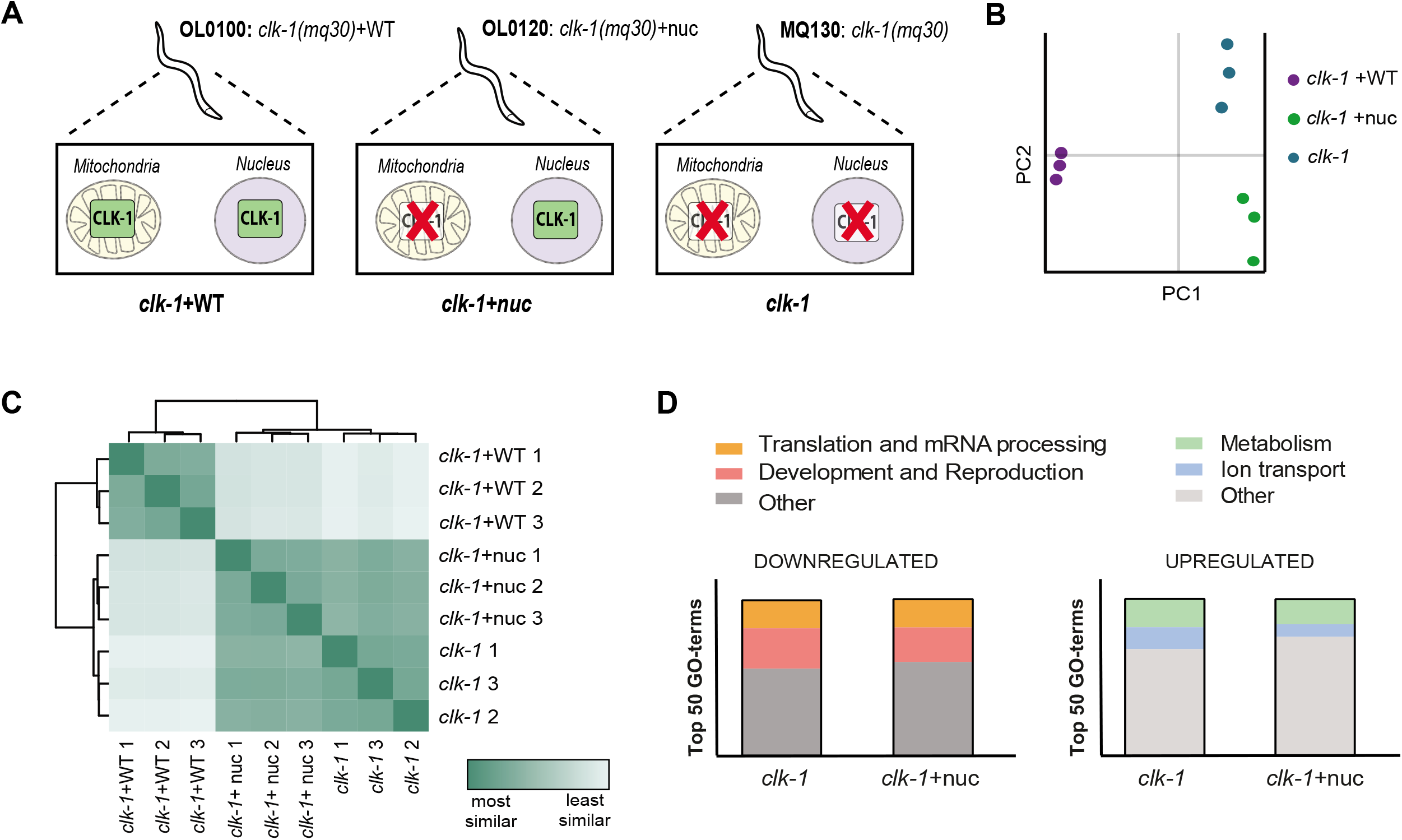
RNA-seq of the *clk-1* (+/-nuclear or WT *clk-1*) mutants. A Graphical illustration of *clk-1* mutant lines used for RNA-seq. B PCA analysis of RNA-seq samples (N = 3 biological replicates for each strain). C Heatmap of RNA-seq samples (N = 3 biological replicates for each strain). D Top 50 downregulated GO-terms (left graph) and upregulated GO-terms (right graph) in clk-1 vs. clk-1 +WT (left column) and clk-1 +nuc vs. clk-1 +WT (right column).

### Transcripts encoding the translation machinery are strongly reduced in *clk-1* strains

We next established the biological pathways involved in *clk-1* mediated longevity using DAVID and GO analysis (Huang da et al., 2009a, Huang da et al., 2009b). Among the most upregulated processes in *clk-1(mq30)* are metabolic pathways (e.g. oxidation-reduction pathway) and ion transport (Fig 1D). A substantial part (25%) of significantly downregulated processes in *clk-1* mutants is involved in reproduction and development (Fig 1D). Developmental delay and reduced progeny have been extensively reported in *clk-1* mutants, and are often described to come at a cost of longevity (Larsen et al., 1995, Tissenbaum & Ruvkun, 1998). Remarkably, another 20% of downregulated GO terms are involved in protein translation and mRNA processing (Fig 1D). In total, 38 genes encoding ribosomal proteins, initiation and elongation factors are downregulated in the *clk-1(mq30)* mutants compared to *clk-1(mq30)^+WT^.* In addition, 28 genes specifically involved in biogenesis and processing of ribosomes were downregulated in the *clk-1(mq30)* worms. Similar downregulation of ribosomal proteins and initiation/elongation factors was observed in the *clk-1(mq30)^+nuc^* mutants. These observations suggest that protein translation is repressed in the *clk-1(mq30)/^+nuc^* mutants, and are in line with the repressed polysome profiles we observed before (Essers et al., 2015).

### There is a minor role for nuclear *clk-1* in transcriptional changes

Since the nuclear form of *clk-1* was suggested to associate with the chromatin in the nucleus (Monaghan et al., 2015), we expected changes in the transcriptome in the mutant expressing exclusively the nuclear form of *clk-1.* However, differentially expressed genes in *clk-1(mq30)^+nuc^* were very similar to the differentially expressed genes *clk-1(mq30)* (Fig 2A). There were 3720 genes similarly upregulated in the *clk-1(mq30)^+nuc^* as in the total *clk-1(mq30)* mutant compared to *clk-1(mq30)^+WT^* and 4397 genes similarly downregulated in both strains compared to the wild-type control (Fig 2B). The 292 upregulated genes and 275 downregulated genes in the *clk-1(mq30)^+nuc^* strain that were not up- or downregulated in the *clk-1(mq30)* strain did not lead to significantly clustered GO terms. The 704 genes upregulated in specifically the *clk-1(mq30)* strain and not in the *clk-1(mq30)^+nuc^* strain were involved in biological processes such as ion transport and oxidation-reduction (Fig 2C). The 805 genes downregulated in the *clk-1(mq30)* strain but not in *clk-1(mq30)^+nuc^* were involved in biological processes such as reproduction and response to DNA damage stimuli (Fig 2C). These processes also appeared when comparing the *clk-1* mutants to wild-type worms.

**Figure 2.**
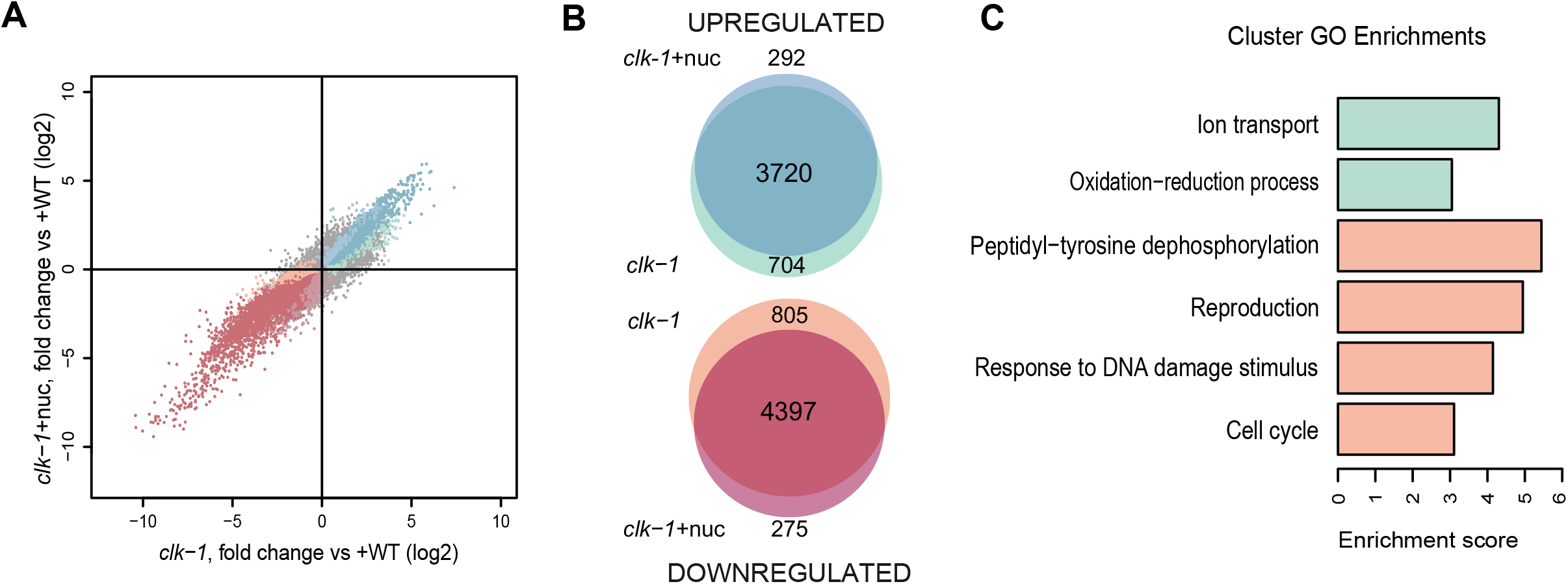
Differentially expressed genes in *clk-1* ^+nuc^ compared to *clk-1*. A Fold changes of differentially expressed genes in *clk-1* vs *clk-1* ^+WT^ plotted against fold changes of *clk-1*^+nuc^ vs *clk-1* ^+WT^. Colors of the individual data points correspond to the colors of the groups of genes in the Venn Diagram in (B). B Venn diagram with number of genes exclusively upregulated (BLUE-GREEN) and downregulated (RED-ORANGE) in either the clk-1^+nuc^ or *clk-1* strain both compared to *clk-1* ^+WT^. Differential expressions in (A) and (B) are with a threshold adjusted *p* value < 0.01. C Significant Cluster GO Enrichments (threshold Enrichment Score > 3) associated with the genes specifically upregulated (GREEN) and downregulated (ORANGE) in *clk-1* strain.

### Translation rates are reduced in *clk-1(mq30)* worms

In order to provide a snapshot of the translational status in the *clk-1(mq30)* strains, we performed polysome profiling of *C. elegans* at the L4 stage. Polysome profiling fractionates total cell lysate over a sucrose gradient by density and enables the distinct measurement of 40S small ribosomal subunits, 60S large ribosomal subunits, 80S monosomes (an mRNA molecule with a single ribosome attached) and multiple polysomes (mRNA molecules with multiple ribosomes attached). Polysome profiles of *clk-1(mq30)* reveal a strong repression of both monosomal and polysomal peaks compared to *clk-1(mq30)^+WT^*, suggesting a global repression of protein translation (Fig 3). Again, the *clk-1(mq30)^+nuc^* mutants show similar yet slightly less dramatic differences compared to wild-type vs. the total *clk-1(mq30)* mutant compared to wild-type (Fig 3). Since the clk-1(mq30)+nuc does not show striking differences in transcription compared to the total clk-1(mq30) mutant, we focused our remaining studies on comparing clk-1(mq30) to clk-1(mq30)+WT. The polysome profiles, together with the RNAseq results, show that *clk-1* mutants are marked by repressed protein translation. Further validation of translational repression in clk-1 mutants is shown by qPCR analysis of several mRNAs encoding EIFs and ribosomal proteins that shows bona fide downregulation of these genes in *clk-1(mq30)* worms (Fig S1).

**Figure 3.**
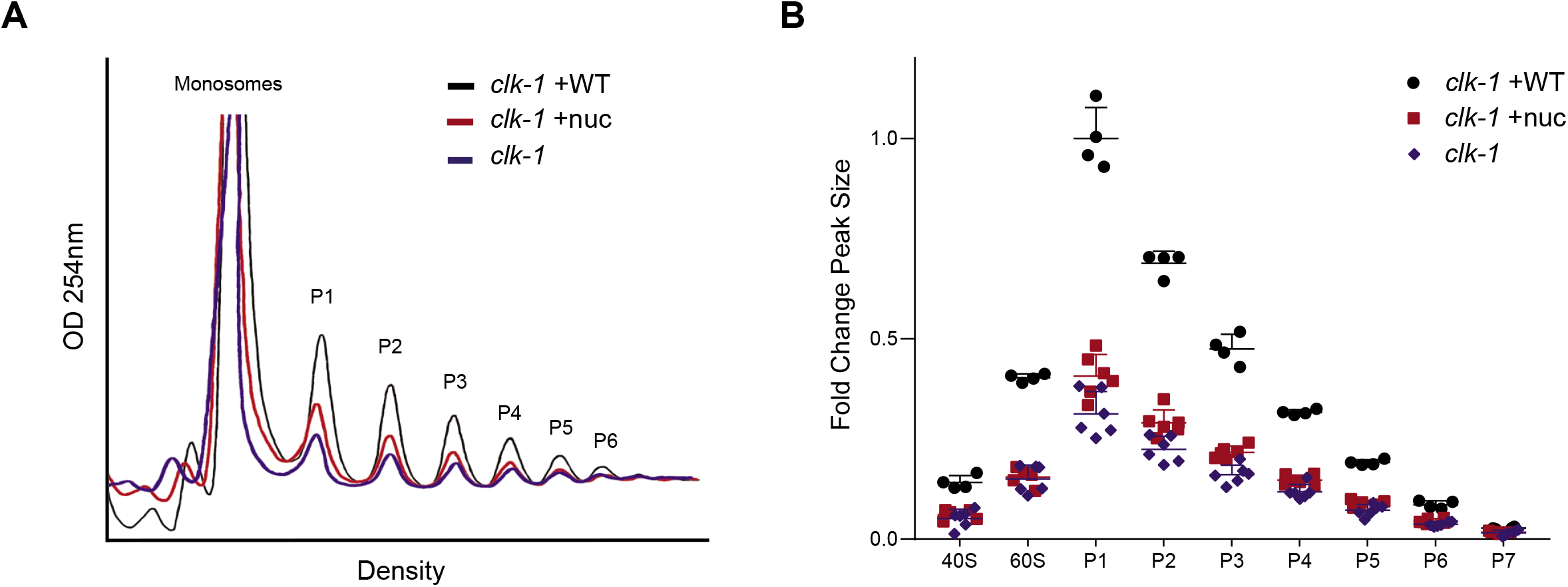
Repressed polysome profiles in the clk-1 (+/-nuc) mutants compared to *clk-1*^+WT^. A Representative polysome profiles of *clk-1* mutants when lysate is normalized to total protein levels of 500ug. The monosomal peak and polysomal peaks are indicated. B) Quantification of polysome peak sizes. The fold change is represented compared to P1 of the clk-1 +WT. All peaks of clk-1+WT are significantly different (P < 0.05) from both *clk-1*+nuc and *clk-1*. No peaks were significantly different between *clk-1* and clk-1+nuc.

### Total RNA does not mirror highly translated polysomal RNA

In addition to providing a global overview of protein translation rates, polysome profiling allows the specific isolation of the RNAs associated with the polyribosomes for comparison with the total RNAs in different *clk-1* strains (Fig 4A). We observed that changes in the polysomal RNA of *clk-1(mq30)* compared to the polysomal RNA of *clk-1(mq30)^+WT^* were not simply mirroring the changes observed in the total mRNA pools of these mutants (Fig 4B). For instance, there are 434 genes specifically upregulated in polysomal RNA pool carrying highly translated messages that are not upregulated in the total RNA pool of *clk-1(mq30)* mutants (Fig 4C). A larger set of 1657 genes is upregulated in both polysomal and total RNA, while the largest part (2767 genes) is exclusively upregulated in the total RNA pool (Fig 4C). A similar pattern is observed with downregulated genes in *clk-1(mq30)* mutants, with 466 genes detected in the polysomal RNA pool, 2574 in both polysomal and total RNA pools, and 2628 in the strictly total RNA pool (Fig 4C)

**Figure 4.**
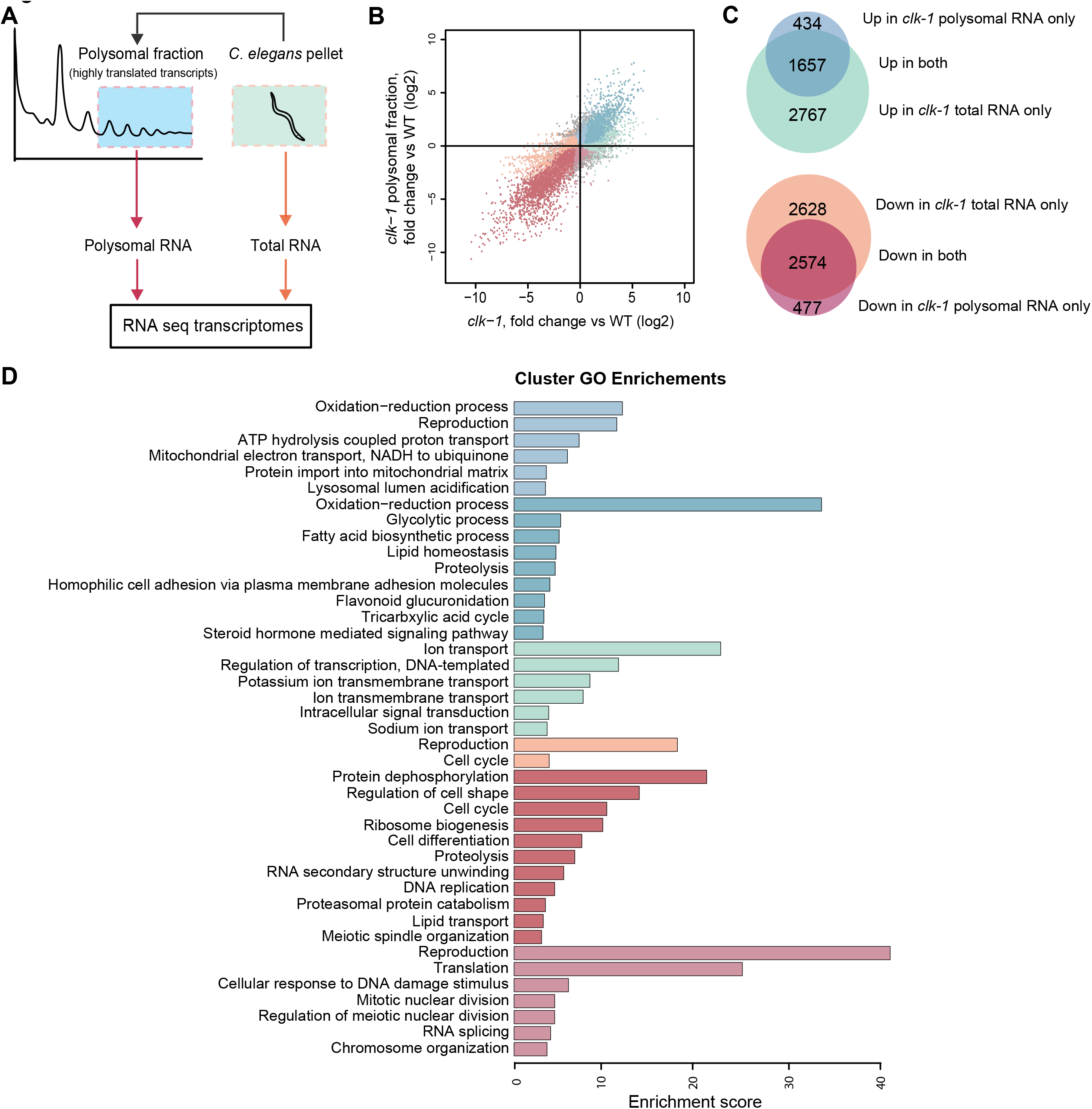
Analysis total and polysomal RNA clk-1 and clk-1+WT. A Schematic representation of polysomal and total RNA from *C. elegans* mutants. B Fold changes of differentially expressed genes in the total RNA of *clk-1* vs *clk-1* ^+WT^ plotted against fold changes in polysomal RNA of *clk-1* vs *clk-1* ^+WT^. Colors of the individual data points correspond to the colors of the groups of genes in the Venn diagram in (C). C Venn diagram with number of genes exclusively upregulated (BLUE-GREEN) and downregulated (RED-ORANGE) in either polysomal RNA or total RNA of *clk-1* strain compared to *clk-1* ^+WT^. Differential expressions in (B) and (C) are with a threshold adjusted *p* value < 0.01. D) Significant Cluster GO Enrichments (threshold Enrichment Score > 3) associated with the genes specifically up and downregulated in groups of genes in (C) (colors of the bars correspond again to the color of the groups in (C).

### OXPHOS transcripts are highly enriched in polysomal RNA of *clk-1(mq30)* strains

In order to determine the functional relevance of the differentially expressed RNAs in total and polysomal RNA pools, we performed DAVID GO analysis. For instance, ion transport processes are upregulated in the total mRNA of *clk-1(mq30)* mutants. However, the polysomal expression data indicate that despite this higher abundance in the total pool, mRNAs coding for ion transport are not more highly translated in *clk-1(mq30)* mutants (Fig 4D). In contrast, mRNAs involved in metabolic processes such as oxidation-reduction, glycolysis, and lipid metabolism are substantially upregulated in both the total and the polysomal RNA pools (Fig 4D). This suggests a metabolic compensation for the reduction of OXPHOS capacity by glycolysis and lipid metabolism in the *clk-1(mq30)* mutants. Processes that were exclusively upregulated in the polysomal RNA pool (and thus exclusively highly translated) in *clk-1* mutants are involved in ATP hydrolysis, protein import into mitochondria, and mitochondrial electron transport (Fig 4D). More specifically, 50 transcripts encoding proteins involved in mitochondrial oxidative phosphorylation were enriched in *clk-1* polysomes (Table 1). This up regulation does not seem complex-specific, since mRNAs coding for proteins of all OXPHOS complexes (I-V) were enriched in *clk-1* polysomes. These data suggest a model whereby proteins involved in ATP production and the import of these proteins into the mitochondrial are highly translated despite lower overall expression levels in *clk-1* mutants compared to WT in order to attempt to overcome their energy deficit.

**Table 1.** Identified polysomal transcripts involved in OXPHOS that, compared with *clk-1^+WT^*, are enriched in *clk-1(mq30)*.

| Accension | Gene name | Description | log2 Fold change<br>Polysomal RNA |
| --- | --- | --- | --- |
| *Complex I* |  |  |  |
| C16A3.5 | C16A3.5 | Orthologue B9 subunit of the mitochondrial complex I | 0.42 |
| C18E9.4 | C18E9.4 | Orthologue B12 subunit of the mitochondrial complex I | 0.61 |
| C25H3.9 | C25H3.9 | Orthologue B5 subunit of the mitochondrial complex I | 0.40 |
| C33A12.1 | C33A12.1 | Orthologue A5/B13 subunit of the mitochondrial complex I | 0.43 |
| C34B2.8 | C34B2.8 | Orthologue A13 subunit of the mitochondrial complex I | 0.48 |
| F37C12.3 | F37C12.3 | Orthologue AB1 subunit of the mitochondrial complex I | 0.53 |
| F42G8.10 | F42G8.10 | Orthologue B11 subunit of the mitochondrial complex I | 0.38 |
| F44G4.2 | F44G4.2 | Orthologue B2 subunit of the mitochondrial complex I | 0.54 |
| F53F4.10 | F53F4.10 | Orthologue NADH-ubiquinone oxidoreductase flavoprotein 2 | 0.45 |
| ZK973.10 | *lpd-5* | NADH-ubiquinone oxidoreductase fe-s protein 4 | 0.37 |
| W10D5.2 | *nduf-7* | NADH-ubiquinone oxidoreductase fe-s protein 7 | 0.47 |
| C09H10.3 | *nuo-1* | NADH Ubiquinone Oxidoreductase 1 | 0.35 |
| T10E9.7 | *nuo-2* | NADH Ubiquinone Oxidoreductase 2 | 0.47 |
| W01A8.4 | *nuo-6* | NADH Ubiquinone Oxidoreductase 6 | 0.47 |
| T20H4.5 | T20H4.5 | Orthologue NADH-Ubiquinone oxidoreductase Fe-S protein 8 | 0.39 |
| Y53G8AL.2 | Y53G8AL.2 | Orthologue A9 subunit of the mitochondrial complex I | 0.48 |
| *Complex II* |  |  |  |
| F42A8.2 | *sdhb-1* | Succinate DeHydrogenase complex subunit B | 0.37 |
| F33A8.5 | *sdhd-1* | Succinate DeHydrogenase complex subunit D | 0.74 |
| *Complex III* |  |  |  |
| C54G4.8 | *cyc-1* | CYtochrome C reductase | 0.35 |
| F42G8.12 | *isp-1* | Rieske iron sulphur protein (ISP) subunit of the mitochondrial complex III | 0.27 |
| R07E4.3 | R07E4.3 | Orthologue subunit VII Ubiquinol-cytochrome c reductase complex III | 0.54 |
| T02H6.11 | T02H6.11 | Ubiquinol-cytochrome c reductase binding protein | 0.40 |
| T27E9.2 | T27E9.2 | Ubiquinol-cytochrome c reductase hinge protein | 0.54 |
| F57B10.14 | *ucr-11* | Ubiquinol-Cytochrome c oxidoReductase complex | 0.75 |
| *Complex IV* |  |  |  |
| F26E4.9 | *cco-1* | Cytochrome C Oxidase | 0.46 |
| Y37D8A.14 | *cco-2* | Cytochrome C Oxidase | 0.46 |
| F26E4.6 | F26E4.6 | Orthologue cytochrome c oxidase subunit 7C | 0.55 |
| F29C4.2 | F29C4.2 | Orthologue cytochrome c oxidase subunit 6C | 0.66 |
| F54D8.2 | *tag-174* | Orthologue cytochrome c oxidase subunit 6A2 | 0.45 |
| Y71H2AM.5 | Y71H2AM.5 | Orthologue cytochrome c oxidase subunit 6B1 | 0.48 |
| *Complex V* |  |  |  |
| C53B7.4 | *asg-2* | ATP Synthase G homolog | 0.79 |
| C06H2.1 | *atp-5* | ATP synthase subunit | 0.45 |
| F32D1.2 | *hpo-18* | Orthologue of ATP synthase, H<sup>+</sup> transporting, mitochondrial F1 complex, ε subunit | 0.56 |
| R04F11.2 | R04F11.2 | Orthologue of ATP synthase, H<sup>+</sup> transporting, mitochondrial Fo complex, ε subunit | 0.51 |
| R53.4 | R53.4 | Mitochondrial ATP synthase subunit f homolog | 0.34 |
| R10E11.8 | *vha-1* | Vacuolar H ATPase 1 | 0.50 |
| R10E11.2 | *vha-2* | Vacuolar H ATPase 2 | 0.63 |
| Y38F2AL.4 | *vha-3* | Vacuolar H ATPase 3 | 0.58 |
| T01H3.1 | *vha-4* | Vacuolar H ATPase 4 | 0.61 |
| VW02B12L.1 | *vha-6* | Vacuolar H ATPase 6 | 0.59 |
| C17H12.14 | *vha-8* | Vacuolar H ATPase 8 | 0.66 |
| ZK970.4 | *vha-9* | Vacuolar H ATPase 9 | 0.43 |
| F46F11.5 | *vha-10* | Vacuolar H ATPase 10 | 0.64 |
| Y38F2AL.3 | *vha-11* | Vacuolar H ATPase 11 | 0.58 |
| F20B6.2 | *vha-12* | Vacuolar H ATPase 12 | 0.34 |
| Y49A3A.2 | *vha-13* | Vacuolar H ATPase 13 | 0.35 |
| F55H2.2 | *vha-14* | Vacuolar H ATPase 14 | 0.49 |
| T14F9.1 | *vha-15* | Vacuolar H ATPase 15 | 0.51 |
| C30F8.2 | *vha-16* | Vacuolar H ATPase 16 | 0.55 |
| Y69A2AR.18 | Y69A2AR.18 | Orthologue of ATP synthase, H<sup>+</sup> transporting, mitochondrial F1 complex, γ subunit | 0.25 |

### RNAs coding for ribosomal proteins are substantially reduced in *clk-1* polysomes

An even more substantial downregulation of transcripts involved in translation and ribosome biogenesis processes was revealed in the polysomal RNA pool of *clk-1* deficient worms compared to *clk-1(mq30)^+WT^*, confirming the repression of translational machinery proteins (Table 2). In the polysomal RNA pool of *clk-1(mq30)* mutants, 62 ribosomal proteins, 16 elongation/initiation factors, 14 amino-acyl tRNA synthetases, and 10 other proteins involved in mRNA processing were downregulated (Table 2). These results are reminiscent of the proteomic analysis demonstrating a significant reduction of ribosomal proteins and translation factors in the *daf-2* insulin receptor mutants (Stout et al., 2013). Furthermore, a subset of these transcripts coding for proteins involved in translation (21 ribosomal proteins, 11 Amino-acyl tRNA synthetases, and 8 other elongation/initiation factors) was even downregulated in the total pool of RNA. This indicates that the downregulation of translation machinery is partially regulated in a transcriptional manner. Taken together, it appears that an attempt to overcome an energy deficit in *clk-1* mutants is coupled to the suppression of ribosome biogenesis and mRNA translation, which would conserve flagging energy for other critical cell processes.

**Table 2.** Identified polysomal transcripts involved in protein translation that, compared with *clk-1^+WT^*, are reduced in *clk-1(mq30)*. Highlighted in grey are the transcript that were upregulated in both the polysomal RNA as the total pool of RNA.

| Accension | Gene name | Description | log2 Fold change |
| --- | --- | --- | --- |
| *Translation* |  |  |  |
| ZC434.5 | *ears-1* | Glutamyl(E) Amino-acyl tRNA Synthetase | -0.51 |
| F31E3.5 | *eef-1A.1* | Eukaryotic translation elongation Factor 1-alpha | -0.67 |
| F25H5.4 | *eef-2* | Eukaryotic translation elongation Factor 2 | -0.53 |
| C27D11.1 | *egl-45* | Eukaryotic translation initiation factor 3 subunit A | -0.50 |
| F11A3.2 | *eif-2Bdelta* | Eukaryotic translation initiation factor 2B subunit Δ | -0.79 |
| Y54E2A.11 | *eif-3.B* | Eukaryotic translation initiation factor 3 subunit B | -0.38 |
| T23D8.4 | *eif-3.C* | Eukaryotic translation initiation factor 3 subunit C | -0.43 |
| R11A8.6 | *iars-1* | Isoleucyl(I) Amino-acyl tRNA Synthetase | -0.26 |
| M110.4 | *ifg-1* | Initiation Factor 4G (eIF4G) family | -0.43 |
| K10C3.5 | K10C3.5 | Othologue of Ria1p | -0.92 |
| R74.1 | *lars-1* | Leucyl(L) Amino-acyl tRNA Synthetase | -0.36 |
| C47E12.1 | *sars-1* | Seryl(S) Amino-acyl tRNA Synthetase | -0.43 |
| F28H1.3 | *aars-2* | Alanyl(A) Amino-acyl tRNA Synthetase | -0.46 |
| K08F11.3 | *cif-1* | COP9/Signalosome and eIF3 complex shared subunit | -0.37 |
| Y41E3.10 | *eef-1B.2* | Eukaryotic translation Elongation Factor | -0.31 |
| Y37E3.10 | *eif-2A* | Eukaryotic Initiation Factor | -1.06 |
| R08D7.3 | *eif-3.D* | Eukaryotic Initiation Factor | -1.61 |
| Y40B1B.5 | *eif-3.J* | Eukaryotic Initiation Factor | -1.24 |
| C17G10.9 | *eif-3.L* | Eukaryotic Initiation Factor | -0.61 |
| C47B2.5 | *eif-6* | Eukaryotic Initiation Factor | -0.53 |
| T05H4.6 | *erfa-1* | Eukaryotic Release Factor homolog | -0.30 |
| F22B5.9 | *fars-3* | Phenylalanyl(F) Amino-acyl tRNA Synthetase | -1.65 |
| T10F2.1 | *gars-1* | Glycyl(G) Amino-acyl tRNA Synthetase | -0.41 |
| F53A2.6 | *ife-1* | Initiation Factor 4E (eIF4E) family | -0.72 |
| B0348.6 | *ife-3* | Initiation Factor 4E (eIF4E) family | -0.55 |
| T05G5.10 | *iff-1* | Initiation Factor Five (eIF-5A) homolog | -0.31 |
| Y54F10BM.2 | *iffb-1* | Initiation Factor Five B (eIF5B) | -0.75 |
| F57B9.6 | *inf-1* | INitiation Factor | -0.31 |
| T02G5.9 | *kars-1* | lysyl(K) Amino-acyl tRNA Synthetase | -0.30 |
| F58B3.5 | *mars-1* | Methionyl(M) Amino-acyl tRNA Synthetase | -0.62 |
| F22D6.3 | *nars-1* | asparaginyl(N) Amino-acyl tRNA Synthetase | -0.64 |
| Y41E3.4 | *qars-1* | glutaminy(Q) Amino-acyl tRNA Synthetase | -0.43 |
| F26F4.10 | *rars-1* | arginyl(R) Amino-acyl tRNA Synthetase | -0.55 |
| C47D12.6 | *tars-1* | Threonyl(T) Amino-acyl tRNA Synthetase | -0.44 |
| Y87G2A.5 | *vrs-2* | Valyl(V) Amino-acyl tRNA synthetase | -0.41 |
| Y80D3A.1 | *wars-1* | tryptophanyl(W) Amino-acyl tRNA Synthetase | -0.67 |
| Y105E8A.19 | *yars-1* | tyrosinyl(Y) Amino-acyl tRNA Synthetase | -0.90 |
| *Ribosome* |  |  |  |
| Y71F9AL.13 | *rpl-1* | Large ribosomal subunit L1 protein | -0.53 |
| B0250.1 | *rpl-2* | Large ribosomal subunit L2 protein | -0.82 |
| F13B10.2 | *rpl-3* | Large ribosomal subunit L3 protein | -0.75 |
| B0041.4 | *rpl-4* | Large ribosomal subunit L4 protein | -0.95 |
| 54C9.5 | *rpl-5* | Large ribosomal subunit L5 protein | -0.58 |
| R151.3 | *rpl-6* | Large ribosomal subunit L6 protein | -0.37 |
| Y24D9A.4 | *rpl-7A* | Large ribosomal subunit L7A protein | -0.44 |
| R13A5.8 | *rpl-9* | Large ribosomal subunit L9 protein | -0.67 |
| JC8.3 | *rpl-12* | Large ribosomal subunit L12 protein | -0.48 |
| C32E8.2 | *rpl-13* | Large ribosomal subunit L13 protein | -0.56 |
| C04F12.4 | *rpl-14* | Large ribosomal subunit L14 protein | -0.30 |
| K11H12.2 | *rpl-15* | Large ribosomal subunit L15 protein | -0.61 |
| M01F1.2 | *rpl-16* | Large ribosomal subunit L16 protein | -0.73 |
| Y48G8AL.8 | *rpl-17* | Large ribosomal subunit L17 protein | -0.45 |
| Y45F10D.12 | *rpl-18* | Large ribosomal subunit L18 protein | -0.66 |
| C09D4.5 | *rpl-19* | Large ribosomal subunit L18A protein | -0.51 |
| E04A4.8 | *rpl-20* | Large ribosomal subunit L20 protein | -0.69 |
| C14B9.7 | *rpl-21* | Large ribosomal subunit L18A protein | -0.51 |
| D1007.12 | *rpl-24.1* | Large ribosomal subunit L20 protein | -0.51 |
| F28C6.7 | *rpl-26* | Large ribosomal subunit L18A protein | -0.29 |
| T24B8.1 | *rpl-32* | Large ribosomal subunit L20 protein | -0.26 |
| F37C12.4 | *rpl-36* | Large ribosomal subunit L18A protein | -0.39 |
| C26F1.9 | *rpl-39* | Large ribosomal subunit L20 protein | -0.98 |
| C09H10.2 | *rpl-41* | Large ribosomal subunit L18A protein | -0.41 |
| Y48B6A.2 | *rpl-43* | Large ribosomal subunit L20 protein | -0.50 |
| B0393.1 | *rps-0* | Small ribosomal subunit S protein | -0.49 |
| F56F3.5 | *rps-1* | Small ribosomal subunit S1 protein | -0.57 |
| C49H3.11 | *rps-2* | Small ribosomal subunit S2 protein | -0.50 |
| C23G10.3 | *rps-3* | Small ribosomal subunit S3 protein | -0.50 |
| Y43B11AR.4 | *rps-4* | Small ribosomal subunit S4 protein | -0.54 |
| T05E11.1 | *rps-5* | Small ribosomal subunit S5 protein | -0.45 |
| Y71A12B.1 | *rps-6* | Small ribosomal subunit S6 protein | -0.83 |
| ZC434.2 | *rps-7* | Small ribosomal subunit S7 protein | -0.39 |
| F40F11.1 | *rps-11* | Small ribosomal subunit S11 protein | -0.57 |
| F37C12.9 | *rps-14* | Small ribosomal subunit S14 protein | -0.51 |
| T08B2.10 | *rps-17* | Small ribosomal subunit S17 protein | -0.41 |
| Y57G11C.16 | *rps-18* | Small ribosomal subunit S18 protein | -0.48 |
| F37C12.11 | *rps-21* | Small ribosomal subunit S21 protein | -0.35 |
| F53A3.3 | *rps-22* | Small ribosomal subunit S22 protein | -0.44 |
| T07A9.11 | *rps-24* | Small ribosomal subunit S24 protein | -0.53 |
| H06I04.4 | *ubl-1* | Small ribosomal subunit S27a protein | -0.37 |
| Y62E10A.1 | *rla-2* | Ribosomal protein, Large subunit, Acidic (P1) | -0.39 |
| F10B5.1 | *rpl-10* | Large ribosomal subunit L10 protein | -0.42 |
| T22F3.4 | *rpl-11.1* | Large ribosomal subunit L11 protein | -1.26 |
| C27A2.2 | *rpl-22* | Large ribosomal subunit L22 protein | -0.47 |
| C03D6.8 | *rpl-24.2* | Large ribosomal subunit L24 protein | -0.68 |
| F52B5.6 | *rpl-25.2* | Large ribosomal subunit L23a protein | -0.82 |
| C53H9.1 | *rpl-27* | Large ribosomal subunit L27 protein | -0.52 |
| R11D1.8 | *rpl-28* | Large ribosomal subunit L28 protein | -0.65 |
| C42C1.14 | *rpl-34* | Large ribosomal subunit L34 protein | -0.53 |
| ZK1010.1 | *rpl-40* | Large ribosomal subunit L40 protein | -0.52 |
| C16A3.9 | *rps-13* | Small ribosomal subunit S13 protein | -0.57 |
| F36A2.6 | *rps-15* | Small ribosomal subunit S15 protein | -0.49 |
| T05F1.3 | *rps-19* | Small ribosomal subunit S19 protein | -0.44 |
| Y105E8A.16 | *rps-20* | Small ribosomal subunit S20 protein | -0.47 |
| F28D1.7 | *rps-23* | Small ribosomal subunit S23 protein | -0.62 |
| Y41D4B.5 | *rps-28* | Small ribosomal subunit S28 protein | -0.64 |
| C26F1.4 | *rps-30* | Small ribosomal subunit S30 protein and ubiquitin | -0.43 |
| F42C5.8 | *rps-8* | Small ribosomal subunit S8 protein | -0.70 |
| F40F8.10 | *rps-9* | Small ribosomal subunit S9 protein | -0.41 |
| W01D2.1 | W01D2.1 | Ortholog of human RPL37 (ribosomal protein L37) | -0.41 |
| Y37E3.8 | Y37E3.8 | Ortholog of human RPL27A (ribosomal protein L27a) | -0.40 |
| *mRNA Processing* |  |  |  |
| W03H9.4 | *cactn-1* | CACTiN (Drosophila cactus interacting protein) homolog | -0.65 |
| F32B6.3 | F32B6.3 | Pre-mRNA processing factor 18 | -0.85 |
| K07C5.6 | K07C5.6 | Homologue of splicing factor SLU7 | -0.95 |
| F33A8.1 | *let-858* | Similarity to eukaryotic initiation factor eIF-4 gamma | -0.47 |
| C04H5.6 | *mog-4* | DEAH helicase | -0.65 |
| EEED8.5 | *mog-5* | DEAH-box helicase 8 | -0.52 |
| C50C3.6 | *prp-8* | Yeast PRP (splicing factor) related | -0.27 |
| Y46G5A.4 | *snrp-200* | Small Nuclear RibonucleoProtein homolog | -0.38 |
| C07E3.1 | *stip-1* | STIP (Septin and Tuftelin Interacting Protein) homologue | -0.84 |
| W04D2.6 | W04D2.6 | Orthologue of human RBM25 | -0.57 |

### Differential translational efficiency (TE) of mRNA reveals a role for TOR signaling

Since there are mRNAs that are selectively being translated in the *clk-1* mutant or the wild-type independently of their abundance in the total mRNA pool, we calculated the translational efficiency (TE) of individual mRNA transcripts by taking the ratio between total RNA and polysomal RNA (Fig 5A). DAVID GO analysis on transcripts with high or low TE revealed similar biological processes as observed in the groups of genes specifically up and downregulated in the polysomal RNA presented in fig 4D (Fig 5B). The list of genes with significantly different TEs was then analyzed using the ‘worm longevity pathway’ in KEGG (Ogata et al., 1999) using pathview (Luo & Brouwer, 2013), to investigate overlap in longevity pathways known in *C. elegans.* Besides hits in the UPR^mt^ and stress resistance, both already described in *clk-1* mutants (Durieux et al., 2011), the results also revealed possible involvement of the mTOR signaling pathway in the *clk-1* mutants. Hence, we also checked the mTOR signaling pathway in KEGG pathview. In *C. elegans*, homologs of TOR pathway genes all extend lifespan in *C. elegans* upon siRNA knockdown (Korta et al., 2012). Interestingly, our RNAseq data show that *let-363* (homologue of TOR) is preferentially translated in wild-type but not in *clk-1* mutants (Fig 5C). Other members of the TOR complex 1 (TORC1) including *daf-15* (RAPTOR), *clk-2* (TEL2), *R10H10.7* (TTI1), also showed decreased TE in *clk-1* mutants (Fig 5C). Moreover, our results show that a key player in autophagy, *lgg-1*, is preferentially translated in *clk-1(mq30)* mutants but not in the *clk-1(mq30)^WT^* (Fig 5C).

**Figure 5.**
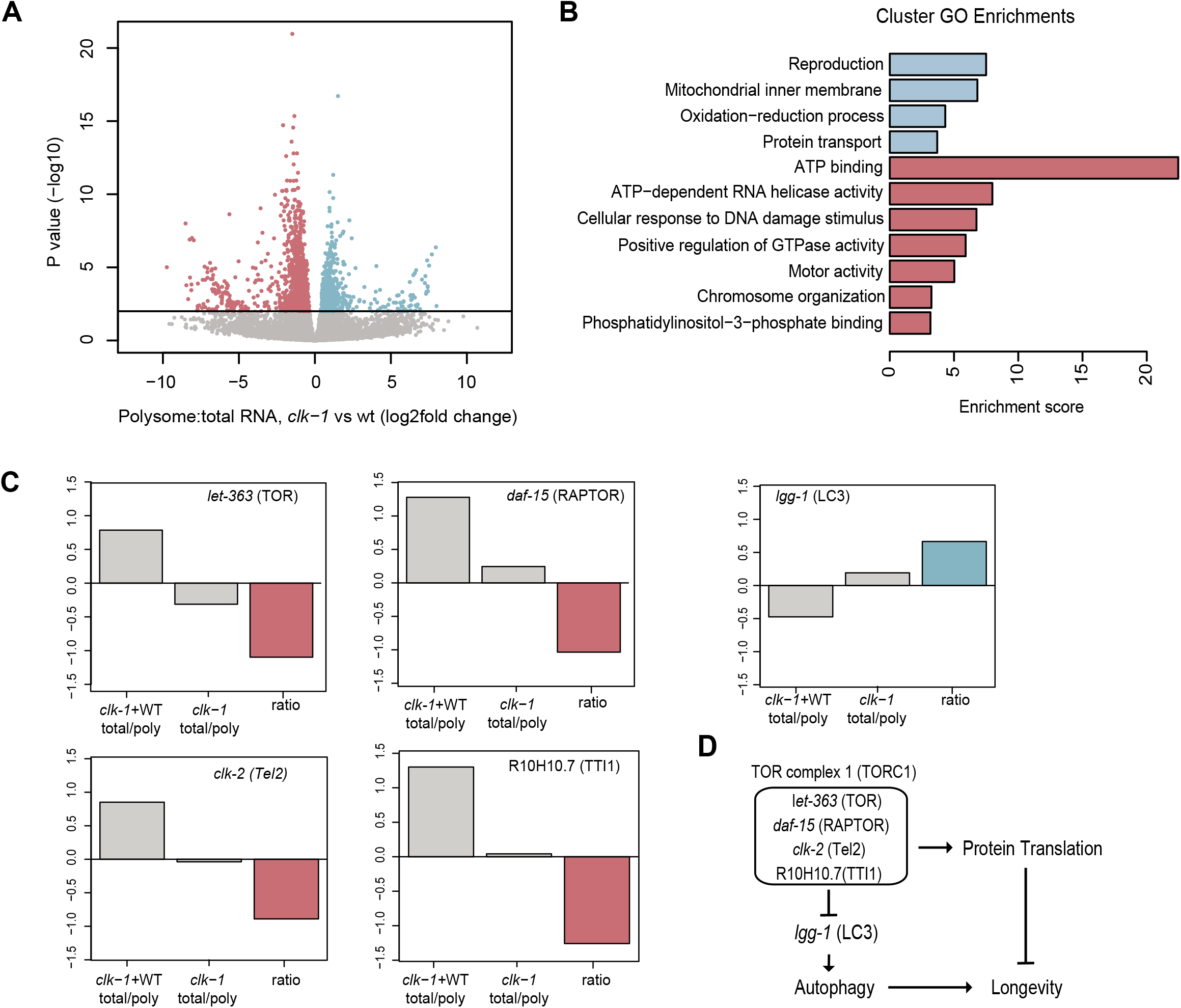
Translation Efficiency (TE) of transcripts in *clk-1* worms. A Volcano plot of Log2 fold-change of TE of transcripts *clk-1*:clk-1+WT. The blue data points represent transcripts with high TE in clk-1 and low TE in clk-1+WT. The red data points represent transcripts that have low TE in *clk-1* and high in *clk-1*+WT. Differentially translationally regulated genes in red and blue with threshold adjusted p value < 0.01. B Significant Cluster GO Enrichments (threshold Enrichment Score > 3) associated with significantly different TEs (colors of the bars correspond again to the data points in (A). C TE of individual transcripts involved in TOR pathway in *clk-1*+wt (left bar) *clk-1* (middle bar) and the ratio (right bar). Colors of the ratio bars correspond again to the data points in (A). D Schematic overview of transcripts involved in TOR pathway that have different translational efficiencies in *clk-1* vs. *clk-1*+WT represented in C and their involvement in longevity.

### *taf-4* is required for repressed polysome profiles in *clk-1(mq30)*

TAF-4 is one of few transcription factors reported to be critical for the *clk-1* longevity phenotype (REF). In order to test if TAF-4 has a role in polysome formation in *clk-1(mq30)* worms, we performed RNAi of *taf-4* in the *clk-1(mq30)* mutants. Here we show that RNAi of *taf-4* significantly increases the monosomal and polysomal peaks of *clk-1 (mq30)* compared to the control empty vector HT115 (Fig 6A,C), suggesting this transcription factor plays a direct role in the suppression of polyribosome formation upon the loss of ubiquinone biosynthesis. This increase is not observed in te polysome profiles upon *taf-4* RNAi in the *clk-1(mq30)^+WT^* (Fig 6B). These data suggest that TAF-4 loss restores polyribosome formation in *clk-1* mutants similar to the previously reported attenuation of the longevity phenotype (Khan et al., 2013). Taken all together, these results confirm a role of reduced cytosolic protein translation in the lifespan extension of the *clk-1* mutants by changing the TE of specific mRNAs coding for the translation machinery and by activating an unappreciated role of TAF-4 in repressing polyribosome formation.

**Figure 6.**
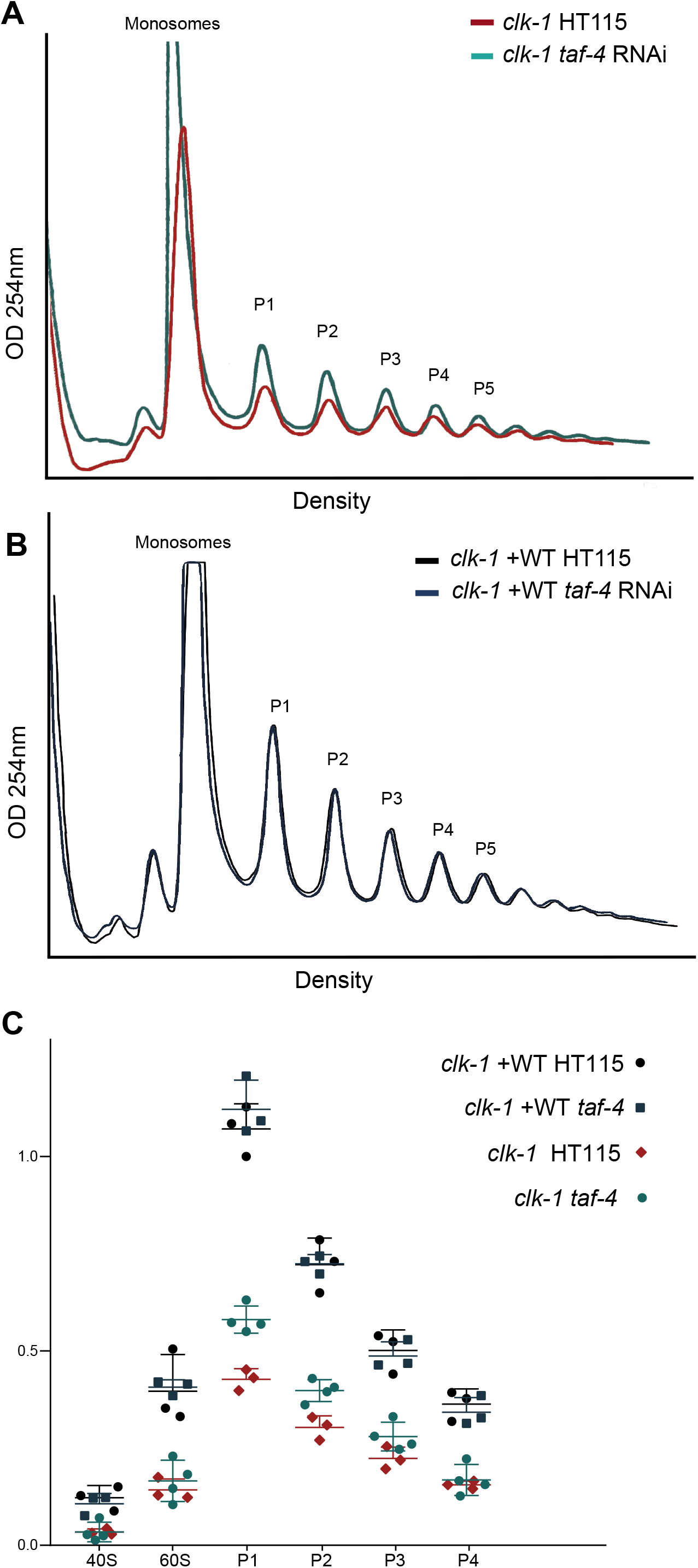
RNAi of *taf-4* partially restores repressed peaks in *clk-1* mutants. A Representative polysome profiles of *clk-1* mutants fed HT115 bacteria transformed with the empty vector or expressing *taf-4* RNAi when lysate is normalized to total protein levels of 500ug. The monosomal peak and polysomal peaks are indicated. B Representative polysome profiles of *clk-1* +WT mutants fed HT115 bacteria transformed with the empty vector or expressing *taf-4* RNAi when lysate is normalized to total protein levels to total protein levels of 500ug. The monosomal peak and polysomal peaks are indicated. C Quantification of polysome peak sizes. The fold change is represented compared to P1 of the *clk-1*+WT controls.

## Discussion

This study was initiated following our previous observation that the polysome profiles of *clk-1* mutants revealed the same repression of polysomes as in the *daf-2* mutants (Essers et al., 2015). We continued this line of inquiry by isolating and sequencing polysomal and total RNAs from *clk-1* mutants and compared them with *clk-1^+WT^* worms. These results demonstrated that compared to *clk-1^+WT^*, very few mRNAs coding for components of the translation machinery such as initiation factors, elongation factors, and ribosomal proteins are present on *clk-1* polyribosomes. In contrast, the *clk-1* polyribosomes reveal a strong preference for translating RNAs coding for OXPHOS proteins. The reduction in the TE of mRNAs coding for the translation machinery is highly reminiscent of the proteomic landscape we described in the *daf-2* mutants, where there is a scarcity of RPs yet also no appreciable reduction in total RP mRNA expression (Depuydt et al., 2013, Essers et al., 2015). Taken all together, the results suggest that shutting down protein translation by altering the TE of mRNAs coding for the translation machinery is a common mechanism used by several different systems that drive lifespan extension.

Our results go on to suggest that the proposed nuclear form of CLK-1 has a relatively small effect on alleviating the polyribosome reduction observed in the *clk-1* mutants. The subtle differences in lifespan and development previously described before (Monaghan et al., 2015) might be explained by reestablishing expression of some reproduction genes in the *clk-1(mq30)^+nuc^* mutant, which are well known to associate to lifespan (Hsin & Kenyon, 1999, Monaghan et al., 2015). Since it has been shown that under certain conditions proteins may be imported into the mitochondria without any MTS (Ruan et al., 2017), it is possible that a small proportion of the nuclear CLK-1 enters the mitochondria without a MTS. This current study does not disprove the existence of nuclear form of CLK-1, however an obvious function of a nuclear CLK-1 is not apparent with these data.

In this study, we confirm the importance of retrograde communication between stressed mitochondria and the rest of the cell. In *Drosophila*, it was shown that dietary restriction-induced longevity, was regulated by activation of the translational repressor 4E-BP, which is the eukaryotic translation initiation factor 4E binding protein (Zid et al., 2009). This resulted in increased ribosomal loading of nuclear-encoded OXPHOS mRNAs. This high TE of mitochondrial proteins was crucial for the dietary restriction-induced lifespan extension, since inhibition of individual mitochondrial subunits of OXPHOS complexes diminished the lifespan extension (Zid et al., 2009). Another study showed that reduction of cytosolic protein synthesis could suppress ageing-related mitochondrial degeneration in yeast (Wang et al., 2008). This postponed mitochondrial degeneration upon reducing cytoplasmic protein synthesis together with our results suggests a robust strategy where mitochondria communicate with the cytoplasmic translation machinery in order to equilibrate cellular energy generated by one compartment and utilized in another. Interestingly, the mitochondrial dysfunction observed in young Mclk1(+/-) mice was shown to paradoxically result in an almost complete protection from the age-dependent loss of mitochondrial function later in life (Lapointe et al., 2009). The increased TE of mitochondrial mRNAs we observed here in the mitochondrial *clk-1(mq30)* mutants could induce this physiological state that ultimately develops long term beneficial effects for aging.

Our results found mRNAs coding for OXPHOS complexes I-V substantially enriched on polysomes of *clk-1(mq30)* worms. Given the marked reduction of total polysomes in these mutants, we can assume that the majority of the few remaining polysomes are present solely to maintain OXPHOS complexes at viable levels. This translational up regulation of OXPHOS mRNAs on ribosomes was also recently observed by ribosome profiling in the context of host shutoff during Vaccinia virus infection in human cells (Dai et al., 2017). This host shutoff facilitates resource availability for viral evasion, and is coupled to a global inhibition of protein synthesis in the host (Bercovich-Kinori et al., 2016). Additionally, the authors propose that this translational up regulation of OXPHOS transcripts is due to their relatively short 5’ untranslated regions (UTRs), which are known to play a regulatory role in TE (Dai et al., 2017) (Araujo et al., 2012). A similar mechanism coupling energy metabolism and translational control is observed in the present study in the context of longevity.

Besides for OXPHOS transcripts, this study also identified an increased TE for the autophagy gene *lgg-1*, the *C. elegans* orthologue of LC3 in humans. Autophagy is an indispensable function in organisms undergoing lifespan extension. In the absence of *bec-1*, the *C. elegans* orthologue of the autophagy gene *APG6/VPS30/beclin1, daf-2* mutant worms fail to undergo lifespan extension (Melendez et al., 2003). Reducing *bec-1* in worms subject to caloric restriction (*eat-2* mutants) or inhibition of the mTOR pathway (reducing let-363 (TOR) or daf-15 (RAPTOR) expression) also has the same effect of preventing lifespan extension (Hansen et al., 2008). Our results suggest a novel mechanism for inducing autophagy in systems undergoing lifespan extension by increasing the TE of mRNAs coding for crucial components of the pathway.

Finally, we found an unexpected role for TAF-4 as a repressor of polyribosome formation when ubiquinone biosynthesis is compromised. Though TAF-4 is essential for all RNA transcription in the early embryo in *C. elegans* (Walker et al., 2001), this requirement is different in later developmental stages and worms exposed to *taf-4* RNAi starting from L1 throughout their remaining life grow normally (Khan et al., 2013). The authors propose a role for TAF-4 in the retrograde response model of longevity in mutants with dysfunctional mitochondria. In this model dysfunctional mitochondria signal via cAMP response element-binding (CREB) proteins to activate TAF-4 in a way that increases life-extending gene expression. Here we show that one of the consequences of TAF-4 loss in *clk-1* mutants, and perhaps also in *isp-1* and *tpk-1* mutants, is a failure of the system that shuts down protein translation in response to mitochondria impairment. This unexpected ability of TAF-4 to control protein translation will be very interesting for further study.

This study demonstrates that molecular pathways of long-lived *clk-1* mutants with dysfunctional mitochondria converge with those of other distinctly different long-lived mutants to reduce cytoplasmic protein translation. It also reveals a striking change in the transcripts that the polyribosomes prefer to translate when mitochondria are compromised. Ribosomal proteins and other proteins part of the translation machinery are very marginally translated while the mitochondrial OXPHOS genes are still robustly translated despite a substantial reduction in the number of polysomes. We propose that this mechanism is likely recapitulated in many more models of lifespan extension, and reaffirms studies that aim to extend human lifespan by reducing protein translation.

## Material and Methods

### Nematode growing and conditions

*C. elegans* strains used: MQ130 *clk-1(qm30)*, OL0100 ukSi1[p*clk-1::clk-1*::gfp, cb-unc-119(+)] and OL0120 ukSi2[p*clk-1::clk-1* ΔMTS (13-187)::gfp, cb-unc-119(+)] were kindly provided by Alan J Whitmarsh and Gino Poulin (Monaghan et al., 2015). Worms were cultured at 20 °C on nematode growth medium (NGM) agar plates seeded with OP50 strain *Escherichia coli*. Synchronized worms were harvested and snap frozen at L4 larval stage for either total mRNA isolation or polysome profiling. For RNAi knock down experiments, synchronized eggs were plated on NGMi (containing 2mM IPTG) with a bacterial lawn of either *E. coli* HT115 (RNAi control strain, containing an empty vector) or *taf-4* RNAi bacteria.

### Polysome Profiling

Gradients of 17-50% sucrose (11 ml) in gradient buffer (110 mM KAc, 20 mM MgAc2 and 10 mM HEPES pH 7.6) were prepared on the day before use in thin-walled, 13.2 ml, polyallomer 14 89 mm centrifuge tubes (Beckman-Coulter, USA). Nematodes were lysed in 500 ml polysome lysis buffer (gradient buffer containing 100 mM KCl, 10 mM MgCl2, 0.1% NP-40, 2 mM DTT and RNaseOUT™ (Thermofischer scientific)) using a dounce homogenizer. The samples were centrifuged at 3500 g for 10 min to remove debris and the supernatant was subjected to BCA protein assay. In all, 500 mg of total protein for each sample was loaded on top off the sucrose gradients. The gradients were ultra-centrifuged for 2 h at 40.000 rpm in a SW41Ti rotor (Beckman-Coulter, USA). The gradients were displaced into a UA6 absorbance reader (Teledyne ISCO, USA) using a syringe pump (Brandel, USA) containing 60% sucrose. Absorbance was recorded at an OD of 254 nm. All steps of this assay were performed at 4°C or on ice and all chemicals came from Sigma-Aldrich (St Louis, MO, USA) unless stated otherwise. Polysome peaks were quantified by measuring area under the curve (AUC) in ImageJ.

### RNA-sequencing

#### Total/Polysomal RNA isolation

For isolation of total RNA, worms were homogenized with a 5 mm steel bead using a TissueLyser II (Qiagen) for 5min at frequency of 30 times/sec in the presence of TRIzol (Invitrogen) and isolation was continued according to manufacturer’s protocol. Polysomal fractions from two experiments were pooled and mRNA was extracted using TRIzol LS (Invitrogen) according to the manufacturer’s protocol. Contaminating genomic DNA was removed using RNase-Free DNase (Qiagen) and samples were cleaned up with the RNeasy MinElute Cleanup Kit (Qiagen) and sequenced by GenomeScan B.V. Leiden (The Netherlands) at a 20M read-depth.

#### Library Preparation

Samples were processed for Illumina using the NEBNext Ultra Directional RNA Library Prep Kit (NEB #E7420) according to manufacturer’s. Briefly, rRNA was depleted from total RNA using the rRNA depletion kit (NEB# E6310). After fragmentation of the rRNA reduced RNA, a cDNA synthesis was performed in order to ligate with the sequencing adapters and PCR amplicification of the resulting product. Quality and yield after sample preparation was measured with the Fragment Analyzer. Size of the resulting products was consistent with the expected size distribution (a broad peak between 300-500 bp). Clustering and DNA sequencing using the Illumina cBot and HiSeq 4000 was performed according to manufacturer’s protocol with a concentration of 3.0 nM of DNA. HiSeq control software HCS v3.4.0, image analysis, base calling, and quality check was performed with the Illumina data analysis pipeline RTA v2.7.7 and Bcl2fastq v2.17.

### Bioinformatics analysis

#### Mapping of the Reads

The quality of the reads in the fastq files was confirmed using FastQC version 0.11.4. (http://www.bioinformatics.babraham.ac.uk/projects/fastqc) FastQC performs multiple quality tests and provides the user with warnings about possible problems with the raw data. Paired end reads were mapped to the *C. elegans* genome using HISAT2 version 2.1.0(Kim et al., 2015). The WBcel235 genome assembly was downloaded from wormbase (http://www.wormbase.org) and used as reference genome for the mapping. After successful mapping, counts per gene were extracted from the. SAM files with HTSeq version 0.9.1(Anders et al., 2015). The - stranded=reverse setting was used, conform the NEB Ultra Directional RNA Library Prep Kit procedure.

#### Statistical Analysis

The statistical analysis was performed in R using DESeq2 version 1.16.1(Love et al., 2014). DESeq2 models the raw counts with the Negative Binomial distribution and normalizes between samples by calculating size factors using the median ratio method (Anders & Huber, 2010).The statistical test is based on a generalized linear model (GLM) using the Negative Binomial model as an error distribution model. To more accurately model dispersion parameters the trend of dispersion to abundance is taken into account using an empirical Bayes procedure. The significance of the GLM coefficients are determined using a Wald test. The p-values are adjusted to correct for multiple testing. In the differential translational regulation analysis the fitted GLM took the following form:

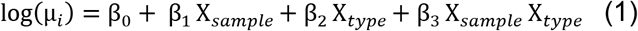

Here the Wald test was performed on the interaction parameter 3 to infer significance. The log2 fold change calculated in this case corresponds to:

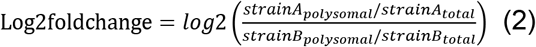

Where in equation (1) sample refers to the strain, and type to polysomal or total RNA. Experimental and statistical normalizations were performed for each condition individually so counts for a specific gene in the polysomal dataset can be possibly higher than in the total RNA dataset, even though the first is technically a subset of the latter. Still, we can test if genes are highly translated in one strain when it is not in the other.

## Acknowledgements

The authors thank Alan J Whitmarsh and Gino Poulin for kindly providing the MQ130, OL0100 and OL0120 *C. elegans clk-1* strains. Work in the Houtkooper group is financially supported by an ERC Starting grant (no. 638290), and a VIDI grant from ZonMw (no. 91715305). Work in the Jelier group is financially supported by a KU Leuven grant C14/16/060. AWM and ML are supported by E-Rare-2, the ERA-Net for Research on Rare Diseases (ZonMW #40-44000-98-1008). GEJ is supported by a 2017 Federation of European Biochemical Society (FEBS) longterm fellowship.

## Author contributions

MM, AWM, and RHH conceived and designed the project. MM and ML performed experiments and GEJ, TS and RJ performed bioinformatics. MM, GEJ, AWM, and RHH interpreted data. MM, AWM and RHH wrote the manuscript, with contributions from all other authors.

---

**Figure S1.**
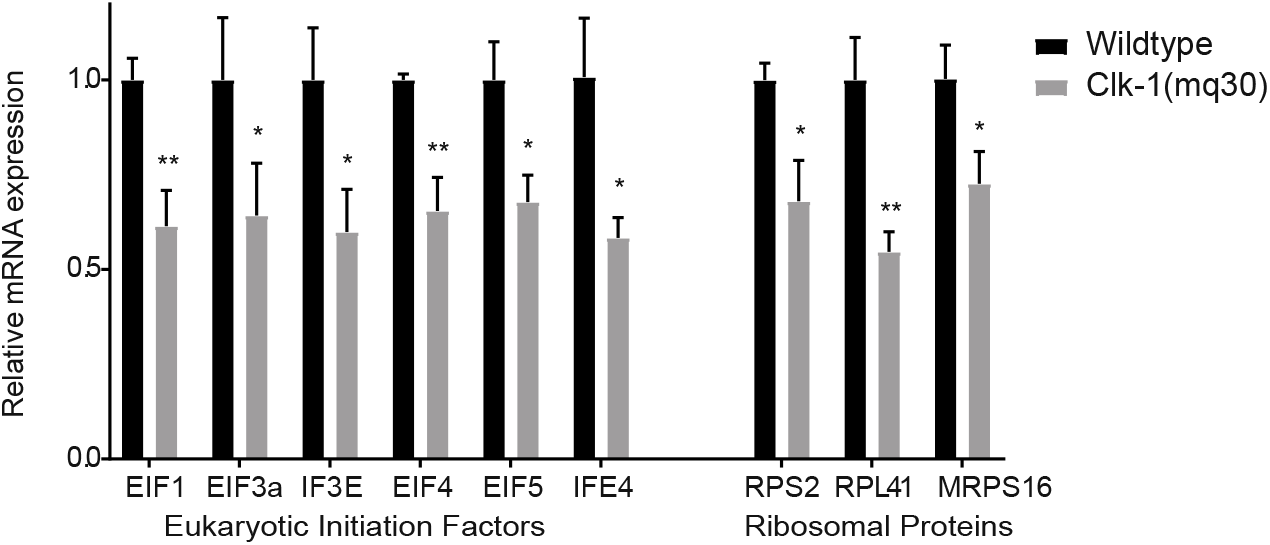
qPCR analysis of EIFs and RPs. Relative expression levels determined by qRT-PCR (Wild-type vs. *clk-1(mq30)*) normalized by expression of pmp-3 as reference gene.

**Figure S2.**
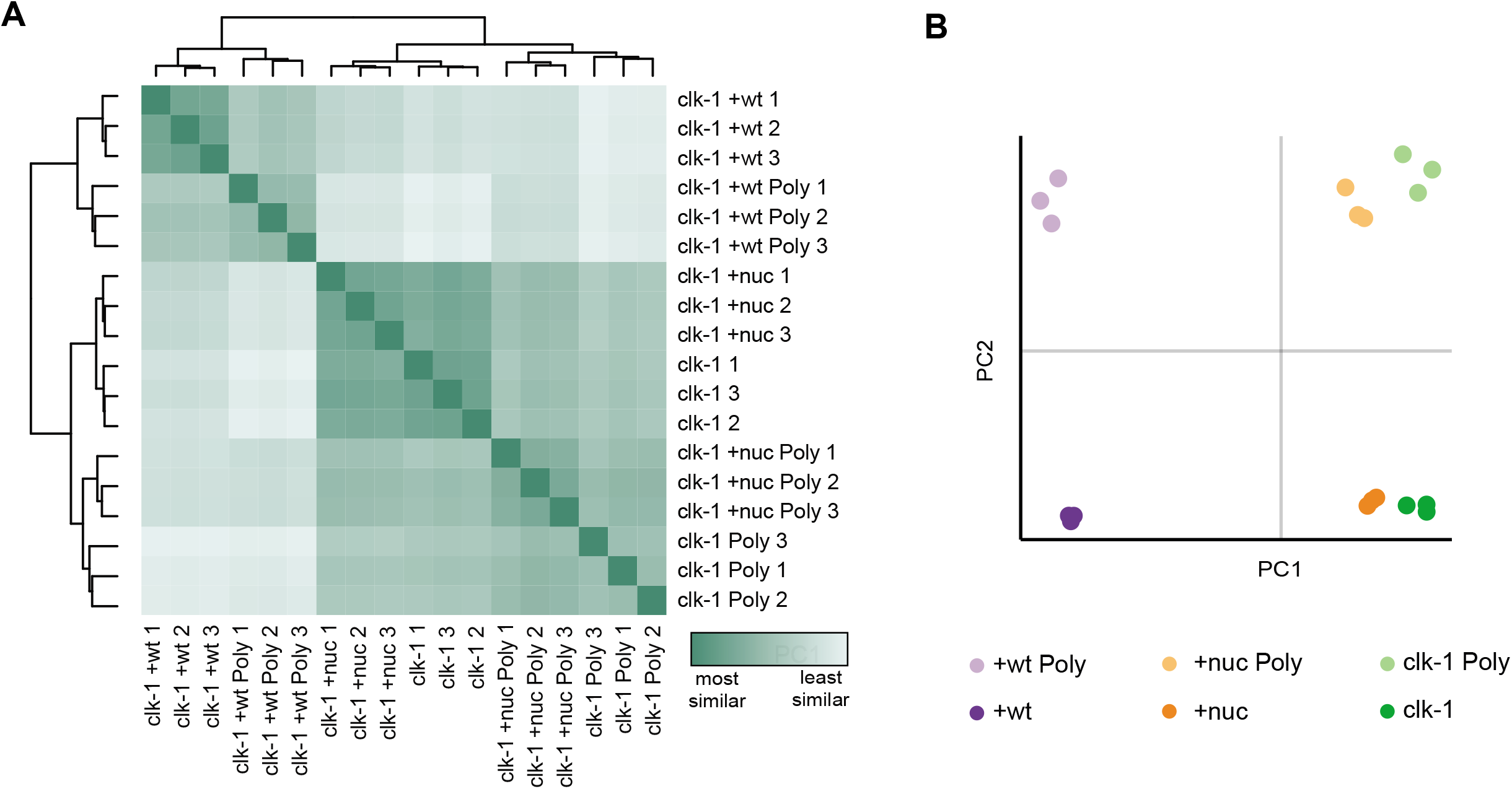
Global analysis of polysomal and total RNA-seq. A Heatmap of (RNA-seq samples (N = 3 biological replicates for each strain). B PCA analysis of RNA-seq samples (N = 3 biological replicates for each strain).

